# Cell fate forecasting: a data assimilation approach to predict epithelial-mesenchymal transition

**DOI:** 10.1101/669713

**Authors:** Mario J. Mendez, Matthew J. Hoffman, Elizabeth M. Cherry, Christopher A. Lemmon, Seth H. Weinberg

## Abstract

Epithelial-mesenchymal transition (EMT) is a fundamental biological process that plays a central role in embryonic development, tissue regeneration, and cancer metastasis. Transforming growth factor-*β* (TGF*β*) is a major and potent inducer of this cellular transition, which is comprised of transitions from an epithelial state to an intermediate or partial EMT state, then to a mesenchymal state. Using computational models to predict state transitions in a specific experiment is inherently difficult for many reasons, including model parameter uncertainty and the error associated with experimental observations. In this study, we demonstrate that a data-assimilation approach using an ensemble Kalman filter, which combines limited noisy observations with predictions from a computational model of TGF*β*-induced EMT, can reconstruct the cell state and predict the timing of state transitions. We used our approach in proof-of-concept “synthetic” *in silico* experiments, in which experimental observations were produced from a known computational model with the addition of noise. We mimic parameter uncertainty in *in vitro* experiments by incorporating model error that shifts the TGF*β* doses associated with the state transitions. We performed synthetic experiments for a wide range of TGF*β* doses to investigate different cell steady state conditions, and we conducted a parameter study varying several properties of the data-assimilation approach, including the time interval between observations, and incorporating multiplicative inflation, a technique to compensate for underestimation of the model uncertainty and mitigate the influence of model error. We find that cell state can be successfully reconstructed in synthetic experiments, even in the setting of model error, when experimental observations are performed at a sufficiently short time interval and incorporate multiplicative inflation. Our study demonstrates a feasible proof-of-concept for a data assimilation approach to forecasting the fate of cells undergoing EMT.

**Author summary:** Epithelial-mesenchymal transition (EMT) is a biological process in which an epithelial cell loses core epithelial-like characteristics, such as tight cell-to-cell adhesion, and gains core mesenchymal-like characteristics, such as an increase in cell motility. EMT is a multistep process, in which the cell undergoes transitions from epithelial state to a partial or intermediate state, and then from a partial state to a mesenchymal state. In this study, we apply data assimilation to improve prediction of these state transitions. Data assimilation is an approach well known in the weather forecasting community, in which experimental observations are iteratively combined with predictions from a dynamical model to provide an improved estimation of both observed and unobserved system states. We show that this data assimilation approach can reconstruct cell state measurements and predict state transition dynamics using noisy observations, while minimizing the error produced by the limitations and imperfections of the dynamical model.

## Introduction

Epithelial-mesenchymal transition (EMT) is a fundamental biological process that plays a central role in embryonic development, tissue regeneration, and cancer metastasis [1–3]. The main characteristic of EMT is the transdifferentiation of an epithelial cell to a mesenchymal cell, which includes losing epithelial-type cell-cell adhesion and gaining the mesenchymal-type enhanced cell motility. While EMT is highly controlled and reversible in embryonic development and wound healing, it is often misregulated in a wide array of disease states, including cancer and fibrotic diseases of the liver, kidney, and heart (reviewed in [1]). In disease states, EMT often progresses unchecked, as opposed to embryonic development, where the process terminates when development is complete. Transforming growth factor-*β* (TGF*β*) is a major and potent inducer of this cellular transition [4–6]. Classically, TGF*β*-induced EMT has been viewed as an all-or-none switch; however, recent work has demonstrated the existence of bistability in the system, with an intermediate or partial state that retains some characteristics of the primary epithelial state but also shows features of the mesenchymal state [7–11]. Thus, we can consider TGF*β*-induced EMT as comprised of transitions from an epithelial state (E) to an intermediate or partial EMT state (P), then to a mesenchymal state (M).

What drives this switch from physiological to pathological EMT? While many of the pathways that drive EMT are understood, the ability to predict when EMT will occur in a reversible process versus when it will proceed unchecked is difficult. The state transition dynamics of the TGF*β*-induced EMT core regulatory pathway is governed by the interactions of a series of transcription factors, including SNAIL 1/2 and ZEB1/2, and their respective inhibition mediated by microRNAs, miR-34 and miR-200 [12]. SNAIL 1/2 and ZEB1/2 induce the state transitions of the epithelial cell by promoting the production of the mesenchymal state marker N-cadherin, while also decreasing the expression of the epithelial state marker E-cadherin. These transcription factors and miRNAs are linked by several feedback loops, including double negative feedback loops between SNAIL1 and miR-34, in which SNAIL1 represses the expression of miR-34 whereas miR-34 negatively regulates the translation of SNAIL1, and between ZEB and its inhibitor miR-200.

Computational modeling of complex cell signaling pathways has become an established tool to understand signaling mechanisms and make predictions of cell states. However, computational models have several key limitations: Even the most detailed biophysical models reproduce the dynamics of a subset of the actual processes occurring in a physiological system. Further, parameters in a computational model are often compiled and extrapolated from a wide range of experimental settings and conditions, and in most cases, parameter values are chosen from experimental mean or median values. While simulations are often valuable tools for understanding mechanisms and interactions between multiple processes with feedback, in general it is difficult to perform computational predictions that correspond with a specific individual experiment. That is, simulations may be representative of the “typical” system behavior, but not reflective of an *individual* experiment. Further, long-term computational predictions will often greatly deviate from the truth, due to even a small degree of uncertainty in parameters and the highly nonlinear nature of biological systems. While there have been efforts to use computational models to generate so-called “populations” of simulations to reproduce inter-trial experimental variability [13, 14], such approaches are generally performed after the experiments to match specific key experimental measurements, and not performed in real time as measurements are made.

Experimental measurements of EMT additionally face the technical challenges associated with direct measurement of multiple epithelial and mesenchymal cell markers. It is not feasible to directly measure all critical EMT-associated cell markers in a given experiment, resulting in incomplete information of cell state. Further, for the cell markers that are measured in a given experiment, calibrating fluorescence intensity measurements and calculating the corresponding expression levels or concentrations of the epithelial- and mesenchymal-associated cell markers is generally not feasible in real time. Ratiometric measurements are one approach to address these calibration issues. Recently, a stable dual-reporter fluorescent sensor was designed to monitor and mirror the dynamic changes of the two key EMT regulatory factors [15], specifically the expression of transcription factor ZEB and epithelial state marker E-cadherin. Importantly, the dual-reporter sensor enables live measurement of the E-cadherin and ZEB ratio. In this study, we demonstrate that we can incorporate these ratiometric measurements (accounting for experimental noise) into a computational approach known as data assimilation, a well-established technique in the atmospheric science field, to accurately forecast cell fate, including predicting *unmeasured* cell marker expression levels and additionally the timing of EMT-associated state transitions.

Data assimilation uses a Bayesian statistical modeling approach to combine high resolution but imperfect dynamical model predictions with sparse, noisy, but repeated experimental observations [16]. More specifically, data assimilation is an iterative algorithm in which a previous state estimate (referred to as the background) is updated based on new observations to produce an improved state estimate (referred to as the analysis), which is the maximum likelihood estimate of the model state. The improved state estimate is then used to produce the initial condition for the dynamical model to predict or forecast the future system state estimate, and the process iteratively repeats.

While data assimilation approaches have been well-utilized in weather forecasting and atmospheric science [16–19], there are relatively few applications in the biomedical sciences [20–30]. In this study, we present a data assimilation approach to reconstruct cell marker expression and predict the timing of the EMT-associated state transitions. We utilize an ensemble Kalman filter, which combines limited noisy observations with predictions from a computational model of TGF*β*-induced EMT [12], to reconstruct the full experimental system and predict the timing of state transitions. We test our approach in proof-of-concept “synthetic” or *in silico* experiments, in which experimental observations are produced from a known computational model with the addition of noise. We mimic parameter uncertainty in *in vitro* experiments by incorporating model error that shifts the TGF*β* doses associated with the state transitions. We find that EMT-associated dynamics can be successfully reconstructed in synthetic experiments, even in the setting of model error, when experimental observations are performed at a sufficiently short time interval. Further, accurate state reconstruction benefits from incorporating multiplicative inflation, a technique to compensate for underestimation of the true background uncertainty (described further below), which helps manage the influence of model error. In summary, our study demonstrates an experimentally feasible data-assimilation approach to cell fate forecasting.

## Methods

The main components of the data assimilation process used in this study are the dynamical systems model (the Tian et al model, described below), the assimilation algorithm (the ensemble Kalman filter), and observations. Here, to establish the validity and accuracy of our approach in an experimental setting, we use synthetic observations, in which the dynamical system is used to generate a known “truth,” with the addition of measurement noise, which can be used for comparison with the data assimilation state estimate. Data assimilation proceeds iteratively, in which simulations of the dynamical model generate a prediction or forecast, after which the ensemble Kalman filter incorporates observations to generate an improved state estimate, known as the analysis step. The improved state estimate then provides the initial conditions of the next forecast (Fig. 1).

**Fig 1.**
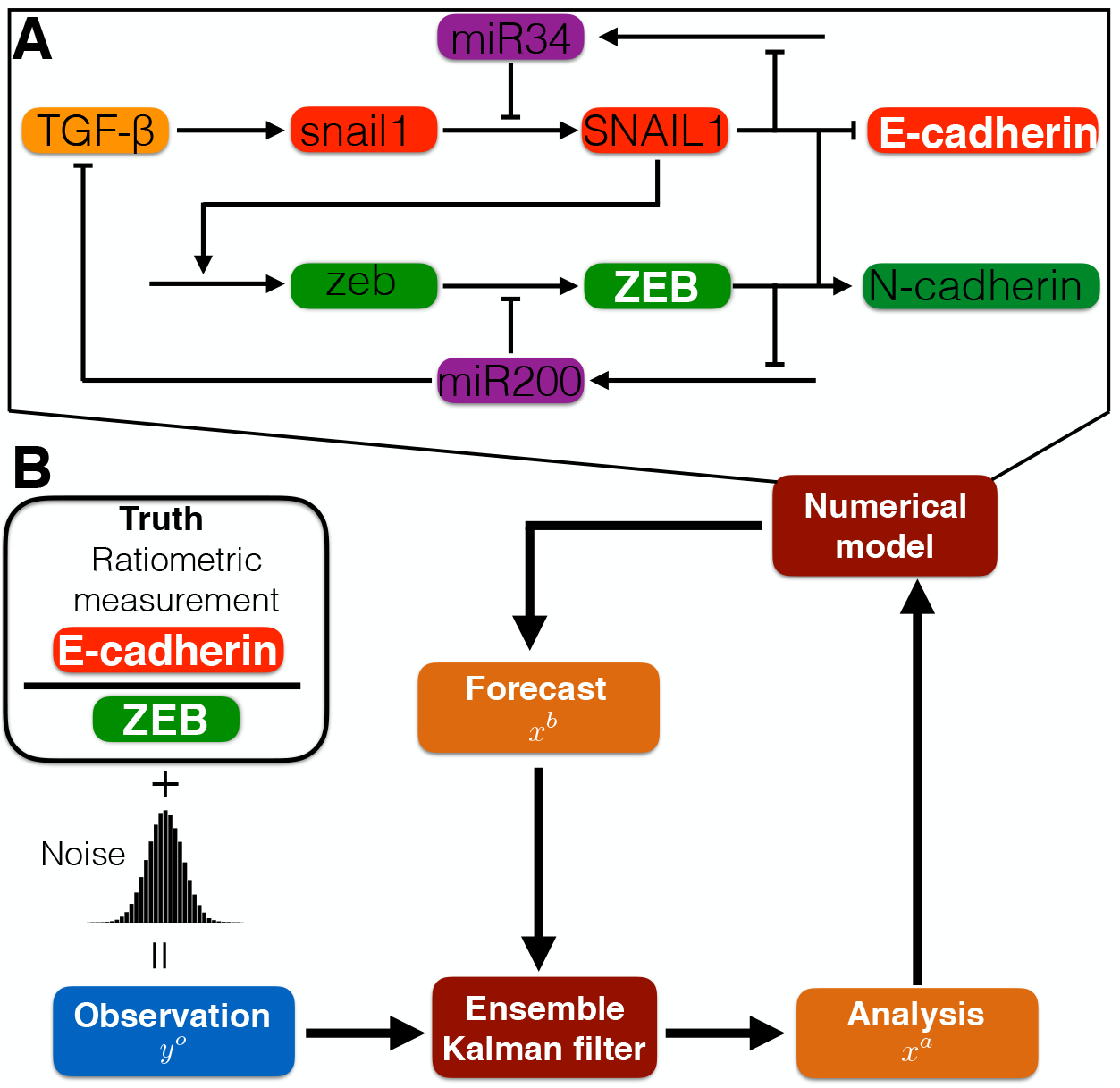
Illustration of EMT regulatory network and data assimilation method. (A) Illustration of the core regulatory network governing EMT dynamics, modified from Tian et al [12] and described in the main text. (B) Diagram of the data assimilation method: Synthetic observations are generated from ratiometric measurements of E-cadherin-to-ZEB from the “truth” system, plus the addition of Gaussian noise. The numerical model (in panel A) generates ensembles of forecasts. Combining the forecasts and observations, the Ensemble Kalman Filter yields the maximum likelihood estimator for the system state (the analysis), which provides initial conditions for the next iteration.

### Computational model of EMT

We use the model from Tian and colleagues to represent the core regulatory network of TGF*β*-induced EMT, given in Eqn. 1 [12]. The dynamics of the system are regulated by two coupled bistable switches, one reversible and the other irreversible. The two bistable switches are regulated by double negative feedback loops, governing the production of transcription factors SNAIL 1/2 and ZEB1/2, respectively, and the inhibition mediated by microRNA miR-34 and miR-200, respectively (Fig. 1A). Model initial conditions and parameters are given in Tables S1 and S2, respectively.

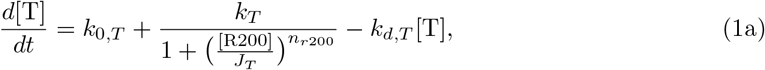

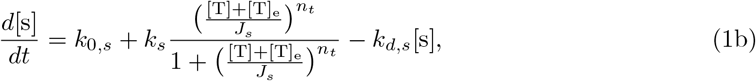

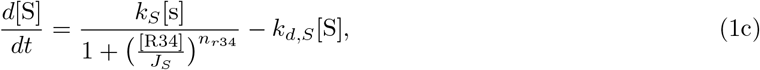

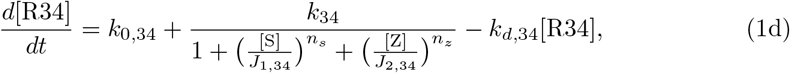

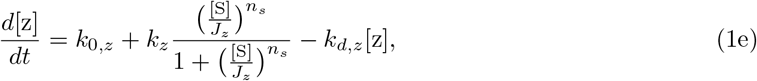

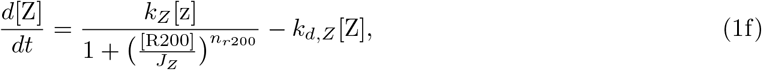

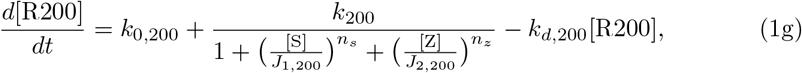

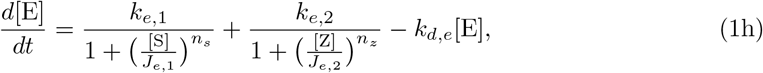

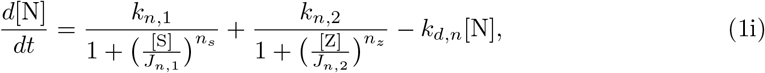

Exogenous TGF*β* increases the production of snail1 mRNA, activating the first double negative feedback loop by increasing the translation of SNAIL1 protein,which in turn inhibits miR-34 production, the inhibitor of SNAIL1 translation. SNAIL1 activates the second double negative feedback loop, by increasing the production of zeb mRNA, increasing translation of ZEB protein, which in turn inhibits miR-200 production, the inhibitor of ZEB translation. Both SNAIL1 and ZEB suppress the epithelial state marker E-cadherin and promote the mesenchymal state marker N-cadherin. Suppression of miR-200 production further removes inhibition of endogenous TGF*β*, a positive feedback that promotes the first feedback loop and results in an irreversible phenotype switch.

A representative simulation demonstrating the transition from an epithelial to mesenchymal state is shown in Figure 2A. Initial conditions are defined consistent with an epithelial state, i.e., high E-cadherin and low N-cadherin expression. A constant dose of 3 *μ*M exogenous TGF*β* is applied for 20 days. An initial increase in SNAIL1 is associated with a moderate decrease in E-cadherin and increase in N-cadherin expression. The simulation illustrates the existence of a state with intermediate levels of both E-cadherin and N-cadherin expression, which is defined as a partial EMT state. A secondary increase of SNAIL1 promotes a subsequent production of ZEB, and in turn, production of endogenous TGF*β*. The transition from the partial EMT to mesenchymal state is associated with a further decrease in E-cadherin and increase in N-cadherin expression.

**Fig 2.**
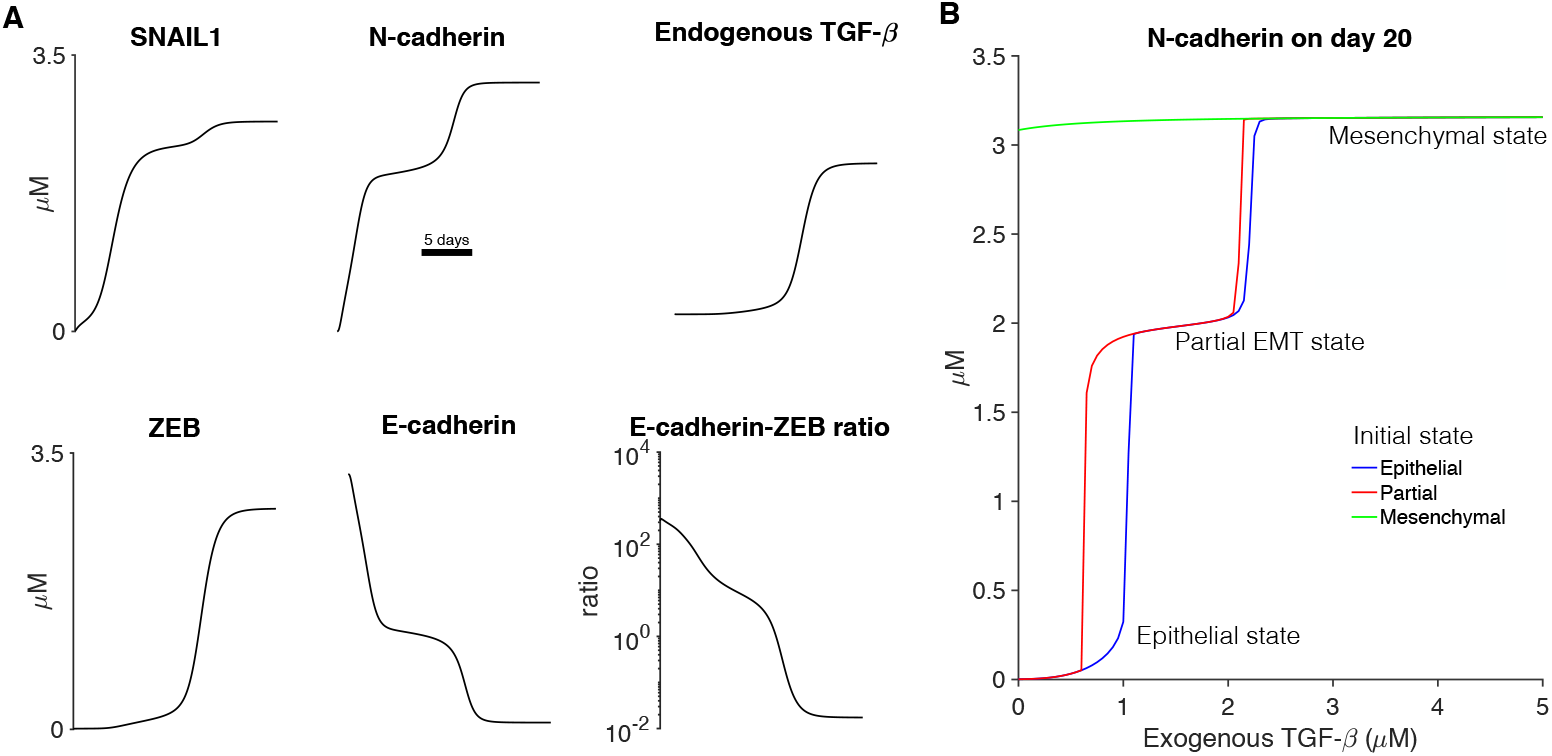
TGF*β* induces EMT via a partial or intermediate EMT state transition. (A) The time course of key epithelial and mesenchymal markers or a ratio of markers are shown as a function, following the addition of exogenous TGF*β*. (B) The expression level of N-cadherin on day 20 are shown as a function of the exogenous TGF*β* dose, for different initial conditions. The step-like response illustrates distinct cell states, corresponding with epithelial, partial or intermediate EMT, and mesenchymal states. Parameters: (A) Exogenous TGF*β* = 3 *μ*M.

Motivated by the recent development of a novel dual reporter sensor for EMT state [15], which emits fluorescence proportional to E-cadherin and ZEB, we also illustrate the dynamics of the ratio between E-cadherin and ZEB, which exhibits a decrease ranging from several orders of magnitudes. As described below, this ratiometric measurement will serve as the observations used in our data assimilation approach, demonstrating the utility of this metric that can be measured experimentally.

In Fig. 2B, we illustrate the model responses to varying exogenous TGF*β* doses for initial conditions in the epithelial (blue), partial (red), or mesenchymal (green) states. We plot N-cadherin expression at the end of a 20-day time interval. For initial conditions in the epithelial state, increasing exogenous TGF*β* results in a step-like increase in the final N-cadherin expression level, with an intermediate level corresponding with the partial EMT state and the elevated level corresponding with the mesenchymal state. Interestingly, for initial conditions in the partial EMT state, hysteresis is observed, such that the TGF*β* doses associated with the epithelial-to-partial state (E-P) transition and the partial-to-mesenchymal state (P-M) transition depend on the initial state. Further, for an initial mesenchymal state, the irreversibility of the second bistable switch results in the maintenance of the mesenchymal state, for all TGF*β* doses, even in the absence of any exogenous TGF*β* added.

### Data assimilation

Data-assimilation methods are a class of algorithms that are used to improve the state-estimation and forecasting ability of dynamical systems by combining observations of the system with a numerical model of the system dynamics. Much of the research on data assimilation for large systems originates in the atmospheric science community [18, 31–33], where it is a crucial piece of numerical weather prediction. While data assimilation was originally designed to improve forecasts by improving the current state estimate—and therefore delaying some of the chaotic drift of the forecast—data assimilation has also been used to estimate and correct parameters of the forecast model. Beyond the Earth’s atmosphere, data assimilation has been used on the Martian atmosphere, as well as on oceans, estuaries, lakes, and biological systems [20, 26, 27, 34, 35].

### Ensemble Kalman filter

In this paper, data assimilation is completed using an ensemble Kalman filter (EnKF), which is an extension of the linear Kalman filter for nonlinear problems. The EnKF attempts to estimate the most likely state of the system given a prior estimate of the state, a (potentially sparse) set of observations of the system, and uncertainty estimates for both the state and the observations. For this problem, the state space at time *t* is a column vector of the nine model variables at this time,

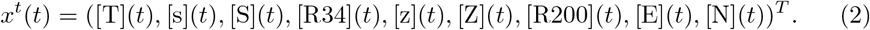

The prior state estimate, which here initially comes from a previous model run, is called the background state and is denoted *x*^*b*^. Estimating the uncertainty in the background—denoted **P**^*b*^—is typically the most difficult piece, especially because this uncertainty is state-dependent. In an EnKF, the background uncertainty is assumed to be Gaussian and the mean and covariance are parameterized by a small number of model states. This is similar to a Monte Carlo approach, but with fewer ensemble members (typically on the order of 10 to 100) than would be needed to fully sample the space.

The algorithm used here is an ensemble transform Kalman filter (ETKF) that is the local ensemble transform Kalman filter (LETKF) algorithm without the localization [16]. Following the notation of Hunt et al. [16], given a set of background states *x*^*b*(*i*)^, the background is computed as the mean of the ensemble members,

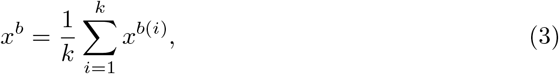

and the covariace is given by the ensemble sample covariance,

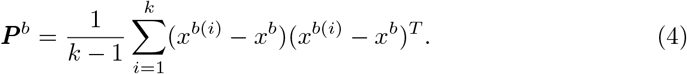

The Kalman filter finds the state that minimizes the cost function

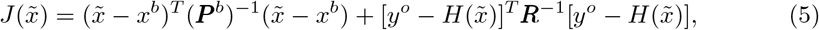

where *y*^*o*^ is the vector of observations, **R** is the covariance of these observations, and *H* is a map from the model space to the observations space (which is typically lower-dimensional). The state that minimizes the cost function in the subspace spanned by the ensemble members is called the analysis and is denoted *x*^*a*^. The analysis error covariance matrix in ensemble space, 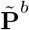, can be computed in ensemble space as 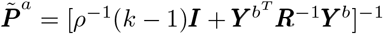. Here, *ρ* is a multiplicative inflation parameter. Multiplicative inflation is a way of compensating for the fact that the small ensemble size tends to lead to underestimation of the true background uncertainty. Multiplying the covariance matrix by a constant greater than 1 (*ρ* here) is the simplest and most computationally efficient way of correcting for this underestimation. The inflation factor, *ρ*, is a tunable parameter for the assimilation. The columns of the **Y**^*b*^ matrix are the perturbations of the background ensemble members mapped into observation space. M athe ically, the *j*th column of **Y**^*b*^ is 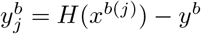, where 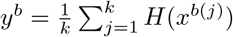 is the mean of the background ensemble in observation space.

The analysis covariance is then used to transform the background ensemble perturbations into analysis ensemble perturbations according to 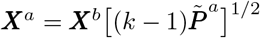. Finally, the new analysis mean is computed as

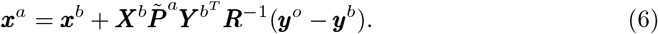

The analysis mean is added to each column of ***X***^*a*^ to generate the analysis ensemble members, which then become initial conditions for the model integration to the next analysis time. Descriptions of the variables in the EnKF method are provided in Table 1. A more detailed description of the algorithm, including derivations, can be found in [16]. In this study, we consider ensemble sizes *k* between 5 and 50, multiplicative inflation factors *ρ* between 1 and 1.6, and observation intervals (i.e., intervals between analysis steps) Δ*t*_*obs*_ between 2 and 48 hours.

**Table 1.** Data assimilation variables. Notation and description of key variables defined and utilized in the Ensemble Kalman filter method for data assimilation. See text for details.

| Notation | Description |
| --- | --- |
| $k$ | Number of ensemble members |
| $m$ | Model space dimension |
| $l$ | Number of observations |
| $\epsilon$ | Gaussian random variable added to the truth to form observations |
| $\rho$ | Multiplicative inflation factor |
| $\Delta t_{obs}$ | Observation interval |
| $H$ | Map from model space to observation space |
| $x^t$ | $m$-dimensional true state vector |
| $x^{b(i)}$ | $m$-dimensional vector of background ensemble member $i$ |
| $y^{b(i)} = H(x^{b(i)})$ | $l$-dimensional vector of the background state estimate mapped to observation space |
| $x^a$ | $m$-dimensional analysis vector |
| $x^b = \frac{1}{k} \sum_{i=1}^k x^{b(i)}$ | $m$-dimensional background state estimate vector |
| $y^b = \frac{1}{k} \sum_{i=1}^k y^{b(i)}$ | $l$-dimensional vector of the mean of $y^{b(i)}$ |
| $y^o = H(x^t) + \epsilon$ | $l$-dimensional observations vector |
| $\mathbf{X}^b$ | $m \times k$ matrix of background ensemble member perturbations from their mean $x^b$ |
| $\mathbf{Y}^b$ | $l \times k$ matrix of background ensemble perturbations in observation space from their mean $y^b$ |
| $\mathbf{X}^a$ | $m \times k$ matrix of analysis ensemble member perturbations from their mean $x^a$ |
| $\mathbf{P}^b$ | $k \times k$ ensemble sample covariance |
| $\mathbf{R}$ | $l \times l$ observation covariance matrix |
| $\tilde{\mathbf{P}}^a$ | $k \times k$ analysis error covariance matrix |

### Numerical experiments

For a given data assimilation trial, the truth system was initialized with all state variables in the epithelial state. To initialize each ensemble member of the background, a separate model simulation was performed, with a random TGF*β* dose (uniformly sampled between the 0 and given dose for that trial), for a random duration (uniformly sampled between 0 and 20 days), and final state variable concentrations were chosen for the ensemble initial state. Synthetic observations were generated from the truth system using a ratiometric measurement of E-cadherin and ZEB. Observational measurement noise or error was reproduced by adding a Gaussian random variable, with standard deviation equal to 10% of the true ratio magnitude, to the true ratio. Minimum E-cadherin and ZEB concentrations were set to 1.1 × 10^−5^ *μ*M, to avoid negative or undefined ratio values.

We assess the accuracy of a given data assimilation trial with two approaches: (1) We calculate the root mean squared deviation (RMSD) between the true system and the average of the analysis ensembles, summing over all state variables, as a function of time:

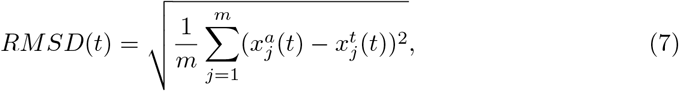

where 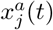 and 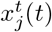 are the *j*th element of the analysis and truth *m*-dimensional vectors, respectively, at time *t*. We calculate the area under the RMSD vs. time curve to quantify error for a single trial. (2) For each ensemble, we predict the timing of the E-P and, where appropriate, P-M state transitions, and compare with the true timing of these transitions. These calculations are performed as follows: After each analysis step, each ensemble is simulated for the remaining time of the 20-day simulation duration. The E-P and P-M state transitions are determined as the time when N-cadherin expression increases above 1.5 and 3.0 *μ*M, respectively. Finally, we average the predicted thresholds over all ensembles. This calculation is repeated for each analysis step.

We first consider the case in which the same parameters are used to simulate both the truth and ensembles, using the baseline parameter set in Tian et al [12]. To assess the data-assimilation approach in the context of parameter uncertainty, we then also consider the influence of model error by increasing the snail1 mRNA degradation rate *k*_*d,s*_ from 0.09 to 0.108. This modification alters the dynamics of the first double negative feedback loop and shifts the TGF*β* doses associated with the E-P and P-M state transitions to higher levels (see Fig. 6). We conducted two variations of our synthetic experiments, with the true system (ensembles) simulated (i) using the baseline (increased) parameter *k*_*d,s*_ value, and (ii) using the increased (baseline) parameter *k*_*d,s*_ value.

Both in the presence and absence of model error, we assessed RMSD and state transition predictions for varying data assimilation properties. Specifically, we varied the time interval between observations/analysis steps Δ*t*_*obs*_, the number of ensembles *k*, and multiplicative inflation *ρ*. For each set of data assimilation properties, measures were averaged over 25 trials to account for randomness in the initialization process.

## Results

A representative data assimilation experiment is shown in Fig. 3, for which the truth (black line) and ensembles (dashed blue lines) utilize the same model parameters (i.e., no model error) and using synthetic E-cadherin-ZEB ratiometric observations (red stars) with an observation interval of 24 hours, 10 ensembles, and no multiplicative inflation (i.e., *ρ* = 1). In both truth and ensemble simulations, 3 *μ*M TGF*β* is applied at time 0. The background ensemble mean (magenta line) followed the true E-cadherin-ZEB ratio within the initial 48 hours (i.e., two analysis steps, Fig. 3B). Importantly, the dynamics of the unobserved state variables, including SNAIL1, ZEB, E-cadherin, N-cadherin, and endogenous TGF*β*, were also reconstructed successfully by the background ensemble mean after 48 hours (Fig. 3A).

**Fig 3.**
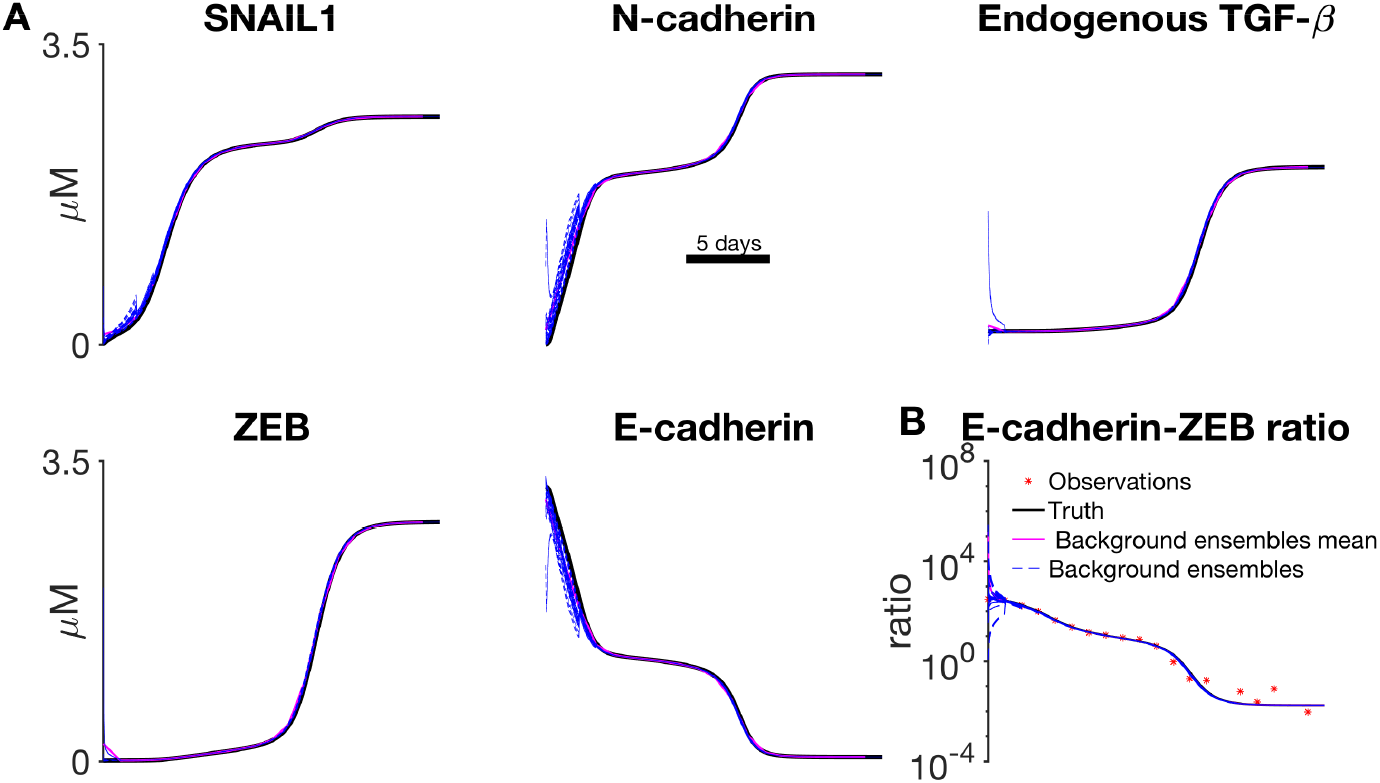
Data assimilation reconstructs unobserved EMT dynamics. (A) The truth (black), ensembles (blue), and ensemble mean (magenta) are shown as a function of time, for key epithelial and mesenchymal markers. (B) Observations (red stars) of the E-cadherin-ZEB ratio are shown for each observation interval Δ*t*_*obs*_. Parameters: Exogenous TGF*β* = 3 *μ*M. Observation interval Δ*t*_*obs*_ = 24 hours, number of ensembles *k* = 10, multiplicative inflation *ρ* = 1.

We first quantified the accuracy of the data-assimilation experiments by measuring RMSD error relative to the true system (Fig. 4A). RMSD error with data assimilation (blue line) demonstrates small increases near the timing of state transitions; however the RMSD error is greatly reduced compared with trials without data assimilation (magenta). We next quantified the accuracy of the data-assimilation corrected simulations to predict the true timing of state transitions from the epithelial-to-partial (E-P) state and partial-to-mesenchymal (P-M) state as follows: After each analysis step, each ensemble was simulated for the remainder of the 20-day duration, and the timing of each transition was determined (if the transition was predicted). We then calculated the mean transition threshold over all ensembles and report this prediction as a function of time (Fig. 4B). We find that the ensemble mean predictions (solid blue lines) initially underestimate the timing of both the E-P and P-M state transitions, due to a subset of ensembles initialized in or near the partial state. However,, the data assimilation predictions converge towards the true timing of E-P and P-M state transitions (black dashed lines) within 48 hours (i.e., two analysis steps), which importantly is before either transition occurs (see Fig. 4B). In contrast, without data assimilation corrections, predictions of both transitions are underestimated (dashed magenta lines).

**Fig 4.**
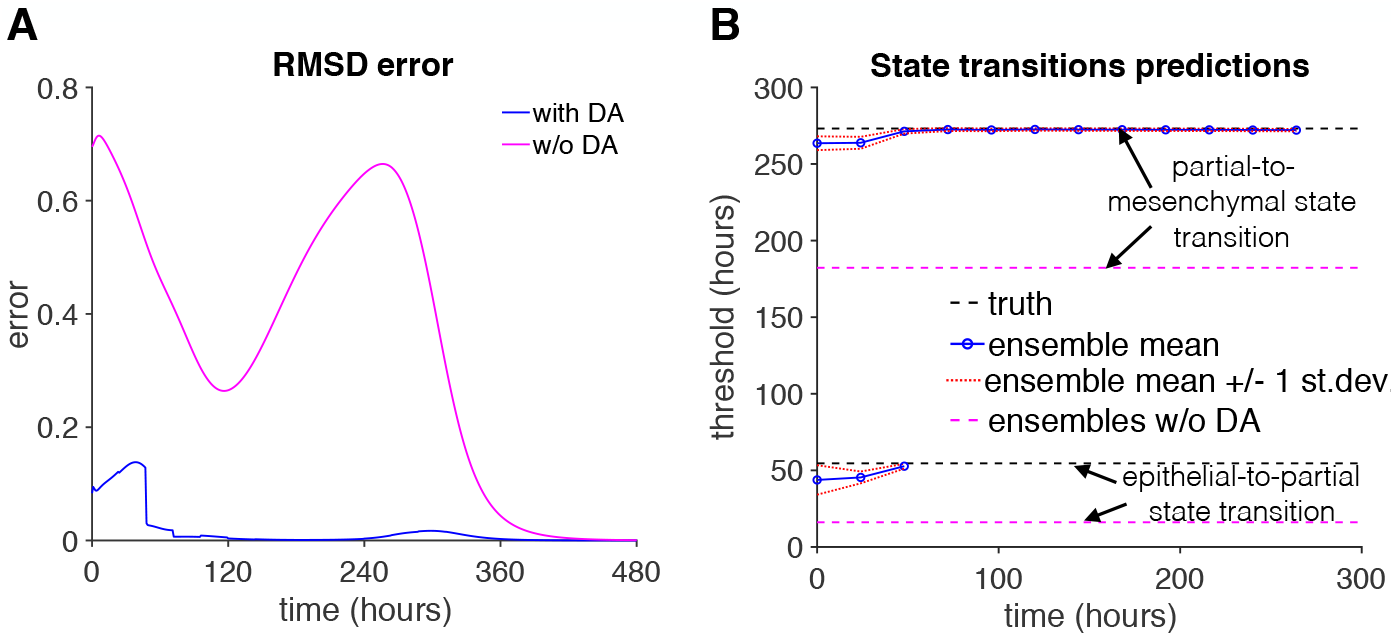
Data assimilation error and predictions. (A) The RMSD error (Eq. 7) for a trial with (blue) and without (magenta) data assimilation (DA) are shown as a function of time. The trial without data assimilation was initialized randomly but no analysis steps were performed. (B) The truth (dashed black) value and ensemble mean (blue), and ensembles without data assimilation mean (dashed magenta) predictions for the epithelial-partial state transition and partial-mesenchymal state transition are shown as a function of time. Ensemble mean predictions ± 1 standard deviation (dashed red) are also shown. Parameters same as Fig. 3.

We next varied the observation interval and number of ensemble members (Fig. 5). For each condition, we calculated the area under the RMSD curve, averaging over 25 trials to account for randomness in the initialization process. Consistent with Fig. 4, for all conditions, RMSD error was much less than the error for trials without data assimilation (dashed magenta). We find that RMSD error area increased approximately linearly as the observation interval increased, i.e., larger error for fewer observations and analysis steps. Interestingly, for these conditions, we also find that varying the number of ensemble members had minimal effect on the RMSD error area, with the exception of a slight increase for 5 ensemble member trials. These results demonstrate that in the absence of model error, this data assimilation approach can greatly reduce error in predictions of system variables and state transition timing, in a manner that depends on the interval for observations while minimally depending on ensemble size. We next consider several conditions in which model error is introduced and further consider key factors that determine the predictive power of the data assimilation approach.

**Fig 5.**
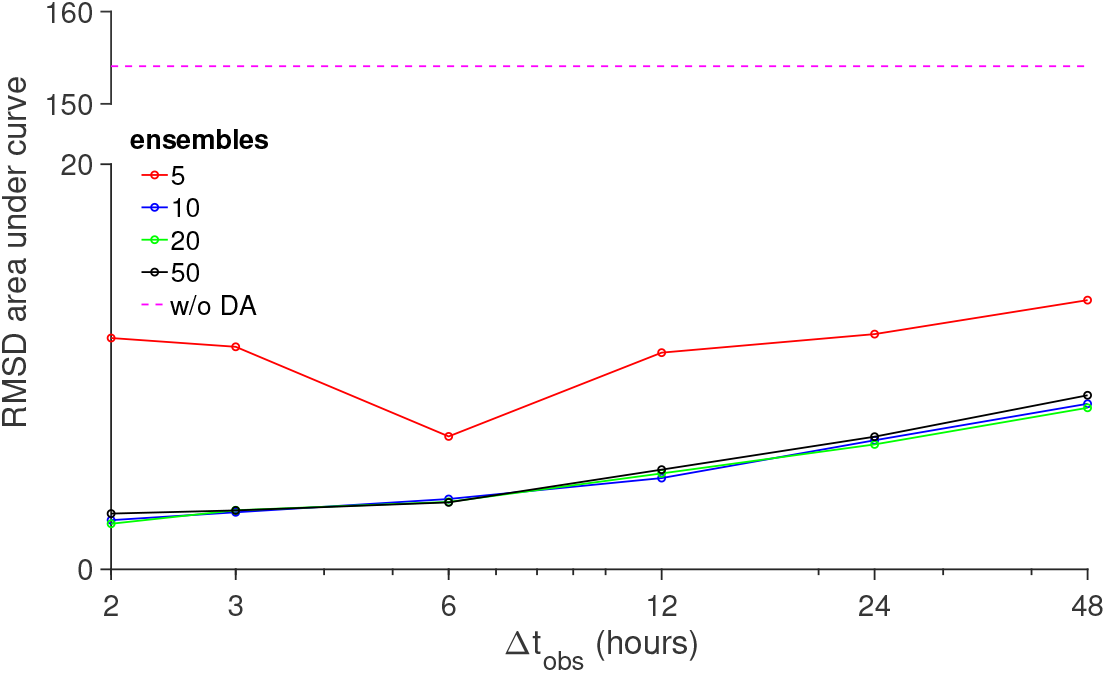
Shorter observation interval reduces prediction error. The area under the RMSD vs time curve is shown as a function of the observation interval Δ*t*_*obs*_, for different number of ensembles *k* (solid lines) and without data assimilation (DA, dashed magenta). Parameters: Multiplicative inflation *ρ* = 1.

**Fig 6.**
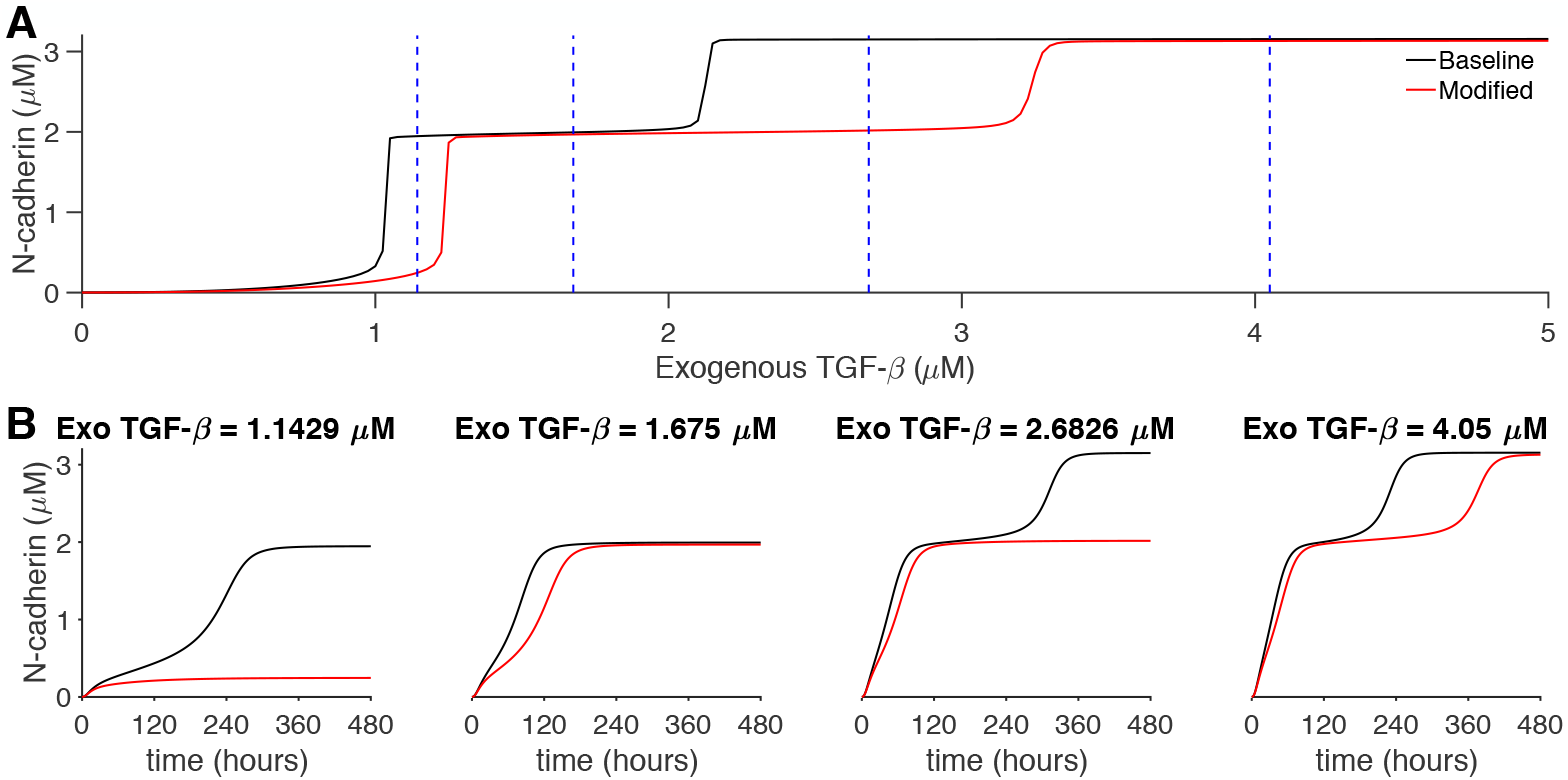
Modified EMT dynamics and TGF*β* dose dependence. (A) N-cadherin expression level on day 20 is shown as a function of the exogenous TGF*β* dose, for baseline (black) and modified (red) parameter sets. (B) N-cadherin expression as a function of time is shown for the four TGF*β* doses denoted in panel A (vertical dashed blue lines). Parameters: baseline: *k*_*d,s*_ = 0.09, modified: *k*_*d,s*_ = 0.108. Other model parameters are unchanged.

We next consider the accuracy of the data assimilation approach in the presence of model error. In particular, we are interested in the situation in which model error results in different steady-state behavior for a given TGF*β* dose. We consider a modified parameter set in which the snail1 mRNA degradation rate *k*_*d,s*_ is increased from 0.09 to 0.108. This modification (red line, Fig. 6A) alters the dynamics of the first double-negative feedback loop and right-shifts the TGF*β*doses associated with the E-P and P-M state transitions to higher levels, relative to the baseline parameter set (black line). Specifically, we consider four exogenous TGF*β* doses test cases (vertical dashed blues lines) that result in four combinations of final steady states between systems with the baseline parameter set and the modified parameter set: (i) 1.1429 *μ*M, which produces a partial EMT state for the baseline system and an epithelial state for the modified system; (ii)1.675 *μ*M, which produces a partial EMT state for both systems, but with altered dynamics; (iii) 2.626 *μ*M, which produces a mesenchymal state on the baseline system and a partial EMT state for the modified system; and (iv) 4.05 *μ*M, which produces a mesenchymal state for both systems, but with altered dynamics (Fig. 6B).

For the next series of synthetic experiments, the true system uses the baseline parameter set, while the ensemble simulations forecast using the modified parameter set with an increased *k*_*d,s*_. Figure 7 illustrates the performance of the data assimilation algorithm for *k* = 20, *ρ* = 1, and Δ*t*_*obs*_ = 6 hours. For exogenous TGF*β* of 1.1429 *μ*M (Dose 1), data assimilation failed to reconstruct the true system dynamics (Fig. 7A, top panel), as the ensemble mean remained in the epithelial state, while the true system transitioned to the partial EMT state. RMSD error was comparable to simulations without data assimilation (middle panel), and furthermore, the E-P transition of the true system was not predicted at any point throughout the simulation (bottom panel).

**Fig 7.**
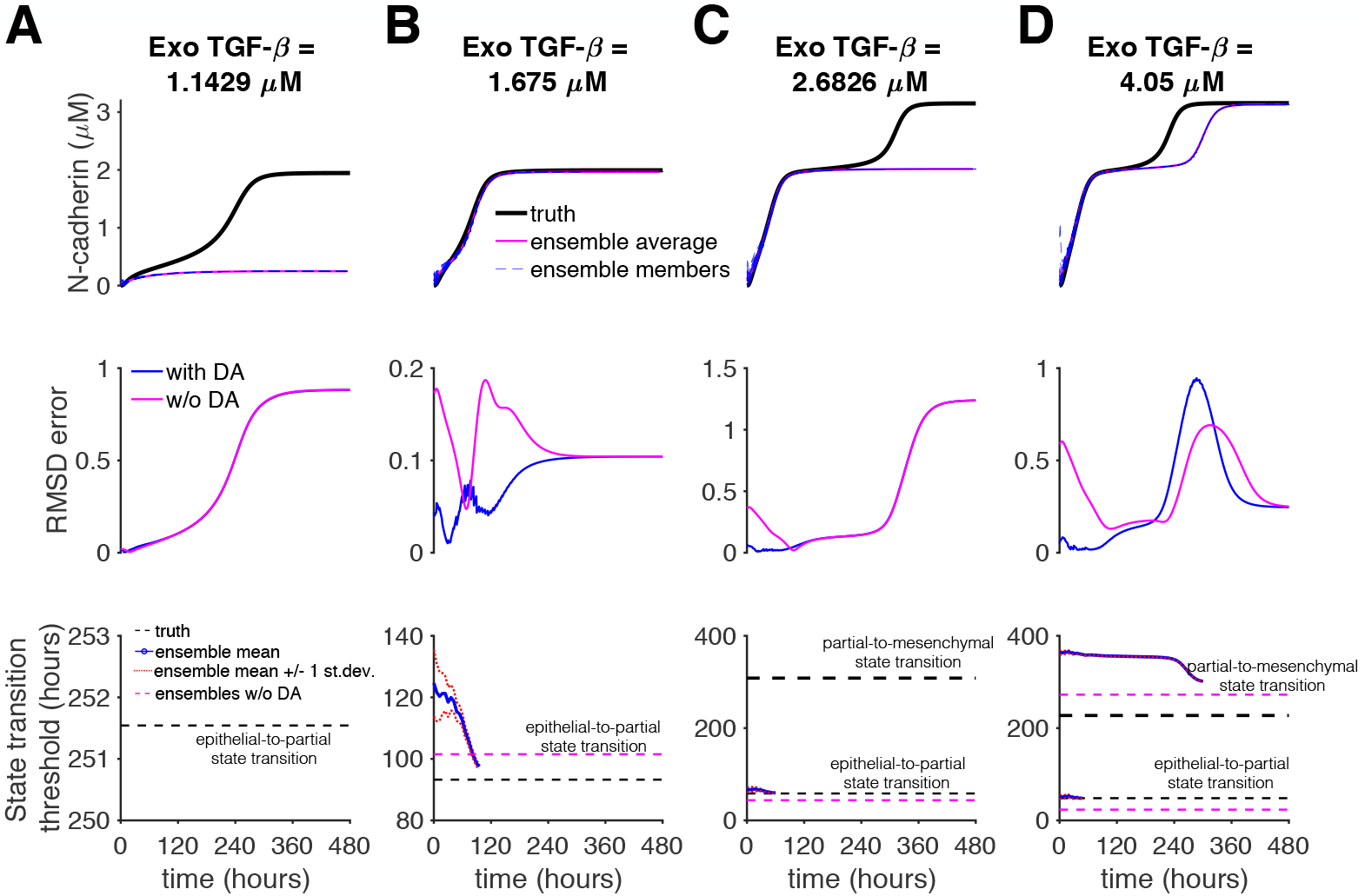
Data assimilation fails to reconstruct EMT dynamics with model error, with frequent observations and without multiplicative inflation. (Top) N-cadherin expression for the truth (black), ensemble members (dashed blue), and ensemble mean (magenta); (Middle) RMSD error with (blue) and without (magenta) data assimilation (DA); and (Bottom) the truth value (dashed black) and ensemble mean (blue) and ensembles without DA mean (dashed magenta) predictions for state transitions are shown as a function of time, for (A-D) four TGF*β* doses. Parameters: observation interval Δ*t*_*obs*_ = 6 hours, number of ensembles *k* = 20, multiplicative inflation *ρ* = 1. Truth system: *k*_*d,s*_ = 0.09, Ensembles: *k*_*d,s*_ = 0.108.

For exogenous TGF*β* of 1.675 μM (Dose 2), data assimilation successfully reconstructed the true system dynamics (Fig. 7B, top panel),with a reduction of the RMSD error before the E-P transition (middle panel). The ensemble prediction of the E-P transition was initially overestimated (consistent with the modified parameter set, see Fig. 6); the prediction improved throughout the simulation, but only accurately predicting the timing immediately preceding the transition (bottom panel).

For exogenous TGF*β* of 2.626 *μ*M (Dose 3), data assimilation failed to reconstruct the true system dynamics, as the ensemble mean remained in the partial EMT state, while the true system transitioned to a mesenchymal state (Fig. 7C, top panel), similar to the first example. Similarly, P-M transition of the true system was not predicted at any point throughout the simulation, although the E-P transition was accurately predicted (bottom panel). Finally, for exogenous TGF*β* of 4.05 *μ*M (Dose 4), data assimilation successfully reconstructed the dynamics of the E-P transition of the true system; the ensemble mean also reproduced the P-M transition although at a later time than the true system (Fig. 7D). The ensemble predictions of both the E-P and P-M transition were initially overestimated; the E-P transition prediction converged on the true timing, while the P-M prediction improved but did not converge before the transition occurred in the true system. Thus, in general, these data assimilation conditions generally failed to predict the timing of state transitions and only predicted steady state behavior when the steady state of true and modified parameter sets were the same (i.e., simulations without data assimilation would also predict the steady state behavior).

We next consider the effect of incorporating multiplicative inflation by increasing *ρ* to 1.4 (Fig. 8). For all TGF*β* doses, the ensemble mean accurately reproduces the dynamics of the true system (top panels) and the RMSD error remained lower than simulations without data assimilation (middle panels). Further, for all exogenous doses, the predictions of the state transitions timings converged to the true values (bottom panels). Importantly, while for TGF*β* Doses 1 and 3, the E-P and P-M transitions, respectively, are initially not predicted to occur, after a sufficient time, these transitions are predicted and indeed converge to the true value (Fig. 8A, C, bottom panels). Thus, we find that incorporating multiplicative inflation greatly improves the predictions of state transitions sufficiently before their respective occurrence in the presence of model error.

**Fig 8.**
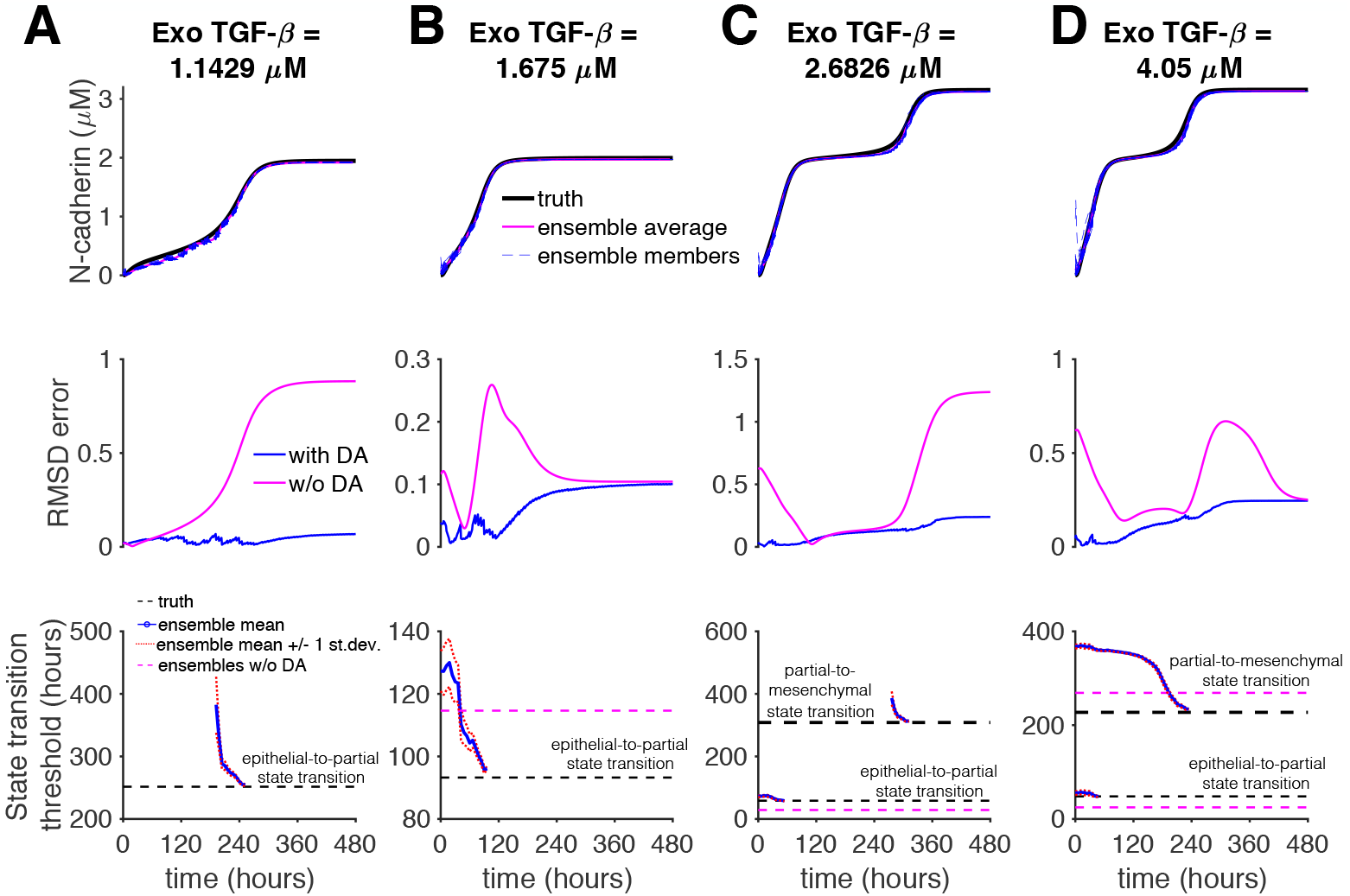
Data assimilation successfully reconstructs EMT dynamics with model error, with frequent observations and multiplicative inflation. (Top) N-cadherin expression for the truth (black), ensemble members (dashed blue), and ensemble mean (magenta); (Middle) RMSD error with (blue) and without (magenta) data assimilation (DA); and (Bottom) the truth value (dashed black) and ensemble mean (blue) and ensembles without DA mean (dashed magenta) predictions for state transitions are shown as a function of time, for (A-D) four TGF*β* doses. Parameters: Observation interval Δ*t*_*obs*_ = 6 hours, number of ensembles *k* = 20, multiplicative inflation *ρ* = 1.4. Truth system: *k*_*d,s*_ = 0.09, Ensembles: *k*_*d,s*_ = 0.108.

With the inclusion of multiplicative inflation, we next investigate the importance of the observation interval by increasing Δ*t*_*obs*_ to 24 hours (Fig. 9). We find that, in general, similar to Fig. 7, the data assimilation approach fails to reproduces the dynamics of the true system. In the case of TGF*β* Dose 1, the steady-state dynamics are predicted, but the timing of the E-P transition is not, while for Dose 3, the steady-state dynamics are not predicted and the P-M transition is not predicted to occur at any point during the simulation. Thus, we find that even with the addition of multiplicative inflation, infrequent observations and analysis step corrections can result in a failure to predict the true system dynamics and associated state transitions.

**Fig 9.**
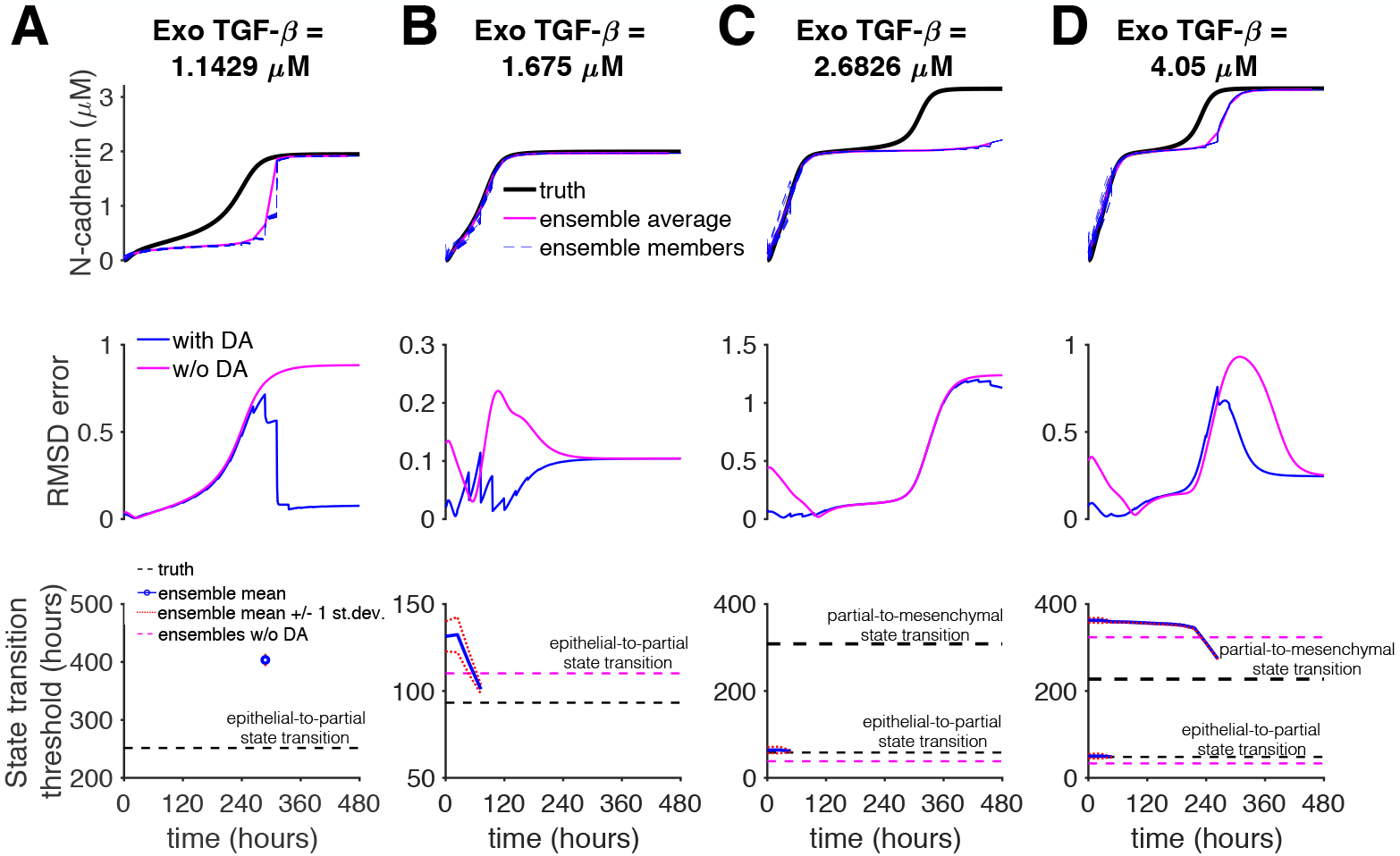
Data assimilation fails to reconstruct EMT dynamics with model error, with infrequent observations and multiplicative inflation. (Top) cadherin expression for the truth (black), ensemble members (dashed blue), and ensemble mean (magenta); (Middle) RMSD error with (blue) and without (magenta) data assimilation (DA); and (Bottom) the truth value (dashed black) and ensemble mean (blue) and ensembles without DA mean (dashed magenta) predictions for state transitions are shown as a function of time, for (A-D) four TGF*β* doses. Parameters: Observation interval Δ*t*_*obs*_ = 24 hours, number of ensembles *k* = 20, multiplicative inflation *ρ* = 1.4. Truth system: *k*_*d,s*_ = 0.09, Ensembles: *k*_*d,s*_ = 0.108.

For this final example, we return to the data assimilation parameters in Fig. 8, with *ρ* = 1.4 and Δ*t*_*obs*_ = 6 hours; however, we consider the case for which the true system used the modified parameter set and the ensembles used the baseline parameters. Similar to Fig. 8, we find that ensemble mean successfully reproduces the dynamics of the true system (top panels), and RMSD error is consistently less than simulations without data assimilation (middle panels). Importantly, for TGF*β* Dose 1, the ensemble mean does not predict the occurrence of the E-P transition. For TGF*β* Dose 3, the P-M transition is initially predicted to occur (consistent with the baseline parameter set, see Fig. 6); however the predicted P-M transition timing gradually increases until the transition is no longer predicted to occur. Finally, for TGF*β* Doses 2 and 4, the timing of the E-P transition and P-M transition (for Dose 4) are initially underestimated and then converge to the true value before the transitions occurred. These simulations thus suggest that data assimilation with properly determined parameters can accurately reproduce the true system dynamics, for conditions in which model error without data assimilation would lead to a failure to predict a state transition (as in Figs. 7-9) or to an erroneous prediction of a state transition (as in Fig. 10).

**Fig 10.**
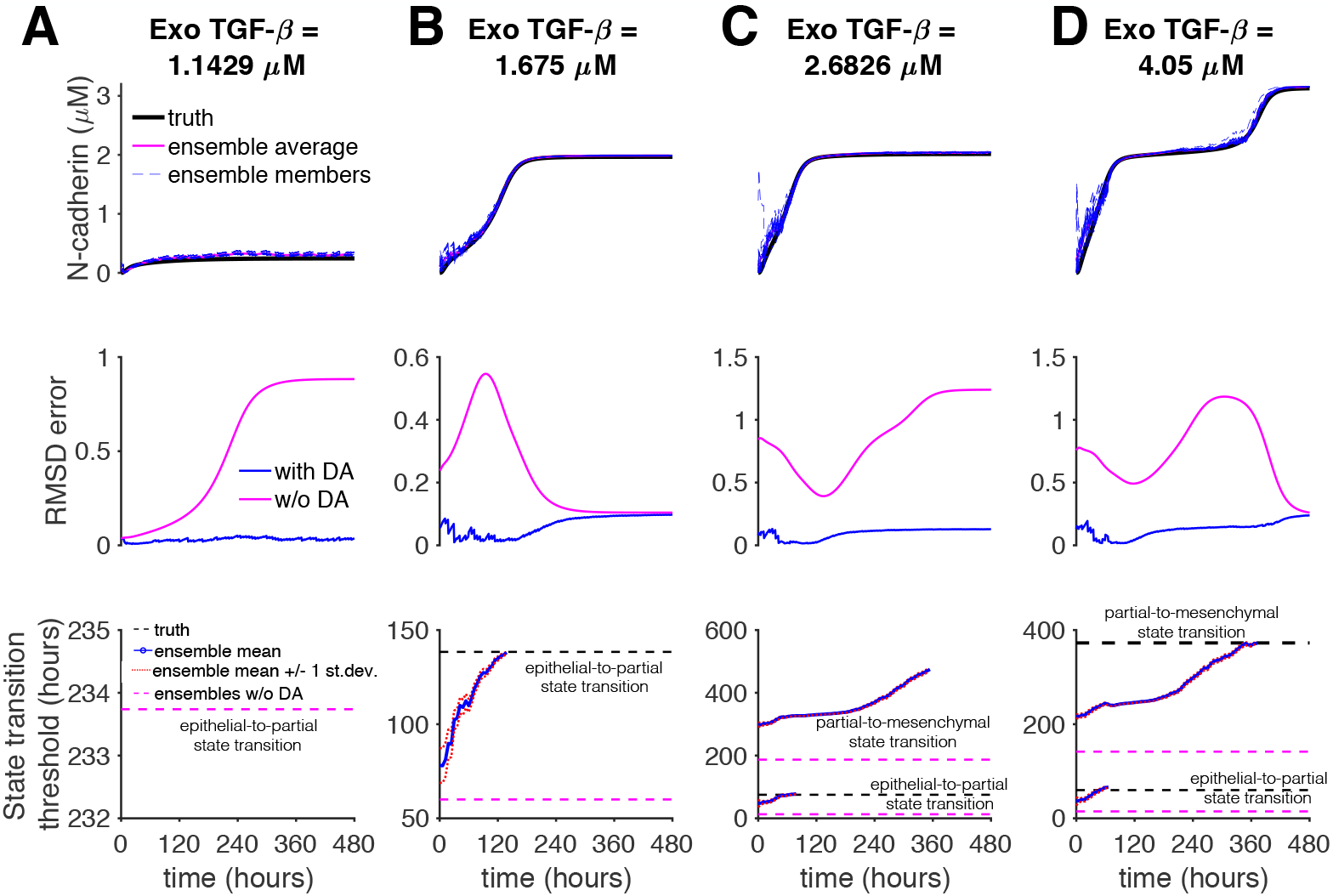
Data assimilation successfully reconstructs modified EMT dynamics with model error, with frequent observations and multiplicative inflation. (Top) N-cadherin expression for the truth (black), ensemble members (dashed blue), and ensemble mean (magenta); (Middle) RMSD error with (blue) and without (magenta) data assimilation (DA); and (Bottom) the truth value (dashed black) and ensemble mean (blue) and ensembles without DA mean (dashed magenta) predictions for state transitions are shown as a function of time, for (A-D) four TGF*β* doses. Parameters: Observation interval Δ*t*_*obs*_ = 6 hours, number of ensembles *k* = 20, multiplicative inflation *ρ* = 1.4. Truth system: *k*_*d,s*_ = 0.108, Ensembles: *k*_*d,s*_ = 0.09.

To broadly quantify the predictive power of the data assimilation approach with the presence of model error, we next performed a parameter study over a wide range of different data assimilation properties, varying *k*, *ρ*, Δ*t*_*obs*_, and the TGF*β* dose. We performed 25 trials for each case and quantified the mean RMSD error area under curve over these trials (Fig. 11).

**Fig 11.**
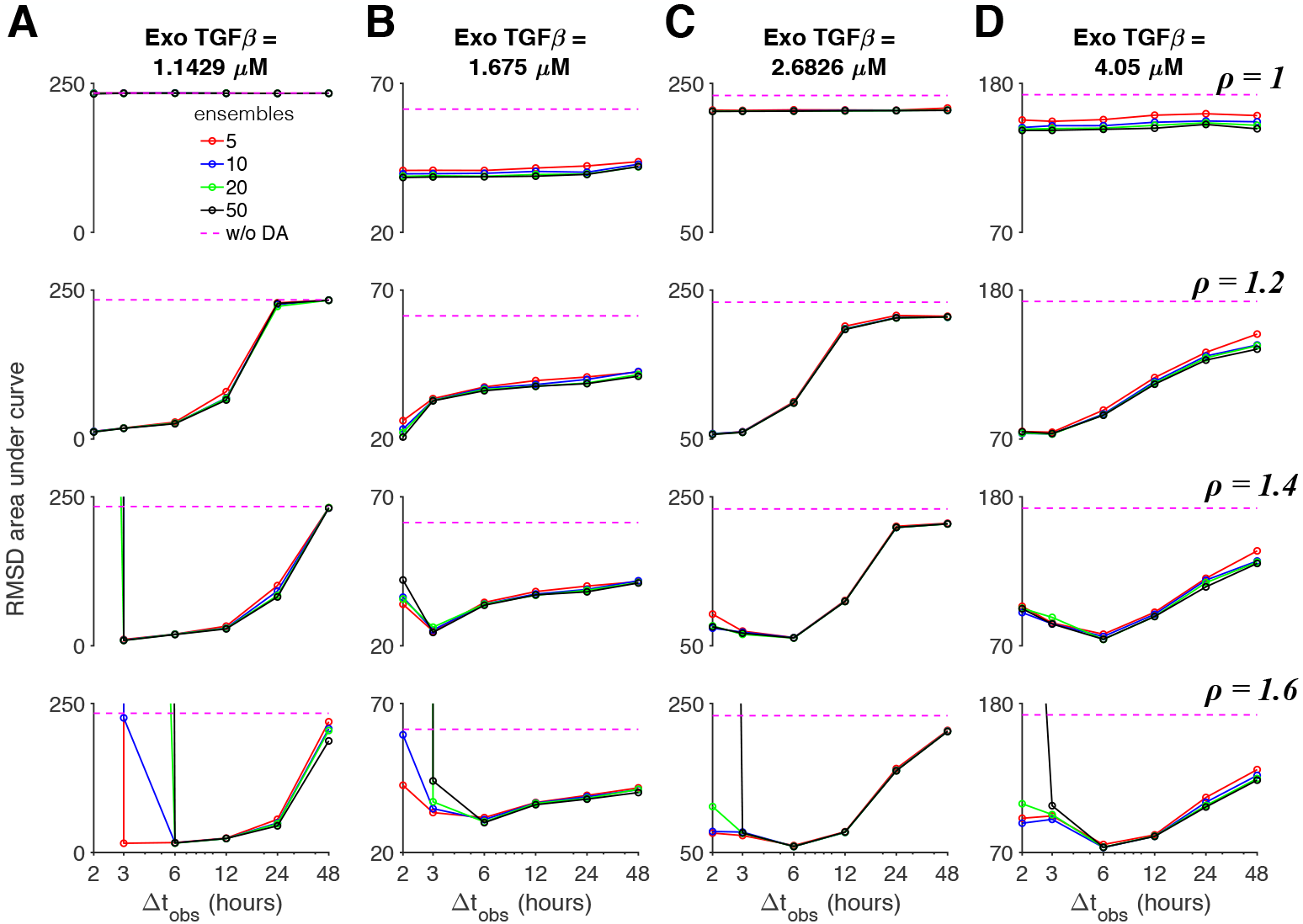
Optimal observation intervals and multiplicative inflation reduce prediction error of EMT dynamics, with model error. The area under the RMSD vs time curve is shown as a function of the observation interval Δ*t*_*obs*_, for different number of ensembles *k* (solid lines) and without data assimilation (DA, dashed magenta), for (A-D) four TGF*β* doses and multiplicative inflation values *ρ* (different rows). Parameters: Truth system: *k*_*d,s*_ = 0.09, Ensembles: *k*_*d,s*_ = 0.108.

For the case of the true system using the baseline parameter set and the ensemble background system using the modified parameter set, we find that in the absence of multiplicative inflation (*ρ* = 1), RMSD error area is only slightly better than simulations without data assimilation (magenta line) for most TGF*β* doses and generally did not depend on observation interval or ensemble size. Consistent with Figs. 7-9, incorporating multiplicative inflation reduced error, generally more so for sufficiently small observation intervals (typically less than 24 or 48 hours). For intermediate multiplicative inflation *ρ* = 1.2, error decreased as the observation interval decreased. For larger multiplicative inflation *ρ* of 1.4 or 1.6, error had a U-shaped dependence, with a minimal error for Δ*t*_*obs*_ of 6 hours typically, for all TGF*β* doses. This demonstrates that while observations are necessary with sufficiently frequent interval to reduce error, too frequent observations (and analysis steps) results in over-correction (i.e., ensemble collapse) and an increase in error. Further, for larger multiplicative inflation (*ρ* = 1.4 − 1.6) and short observation intervals Δ*t*_*obs*_ below 6 hours, a subset of data assimilation trials became unstable, due to state variables in the non-physiological regime, resulting in a dramatic increase in the mean RMSD error, which occurred more frequently for larger ensemble sizes. In general, ensemble size has negligible effect on the error for most cases.

Finally, we performed the same broad data assimilation parameter study, for which the true system used the modified parameter set and the ensemble background the baseline parameter set (as in Fig. 10). Similar to the previous study (Fig. 11), without multiplicative inflation, RMSD error generally does not depend on the ensemble size or observation interval, although for this case, is generally lower than simulations without data assimilation (Fig. 12). Also, as in Fig. 11, error decreases for smaller observation interval Δ*t*_*obs*_ for moderate multiplicative inflation (*ρ* = 1.2) while error generally has a U-shaped dependence for larger multiplicative inflation (*ρ* = 1.4 − 1.6), with a minimum near Δ*t*_*obs*_ of 3 or 6 hours. We similarly find dramatic increases in error for larger *ρ* and small Δ*t*_*obs*_. Thus, across a wide range of data assimilation experiments incorporating model error and multiple TGF*β* doses resulting in different EMT states, we find that moderate multiplicative inflation and short observation intervals consistently demonstrate the lowest state variable predictive error.

**Fig 12.**
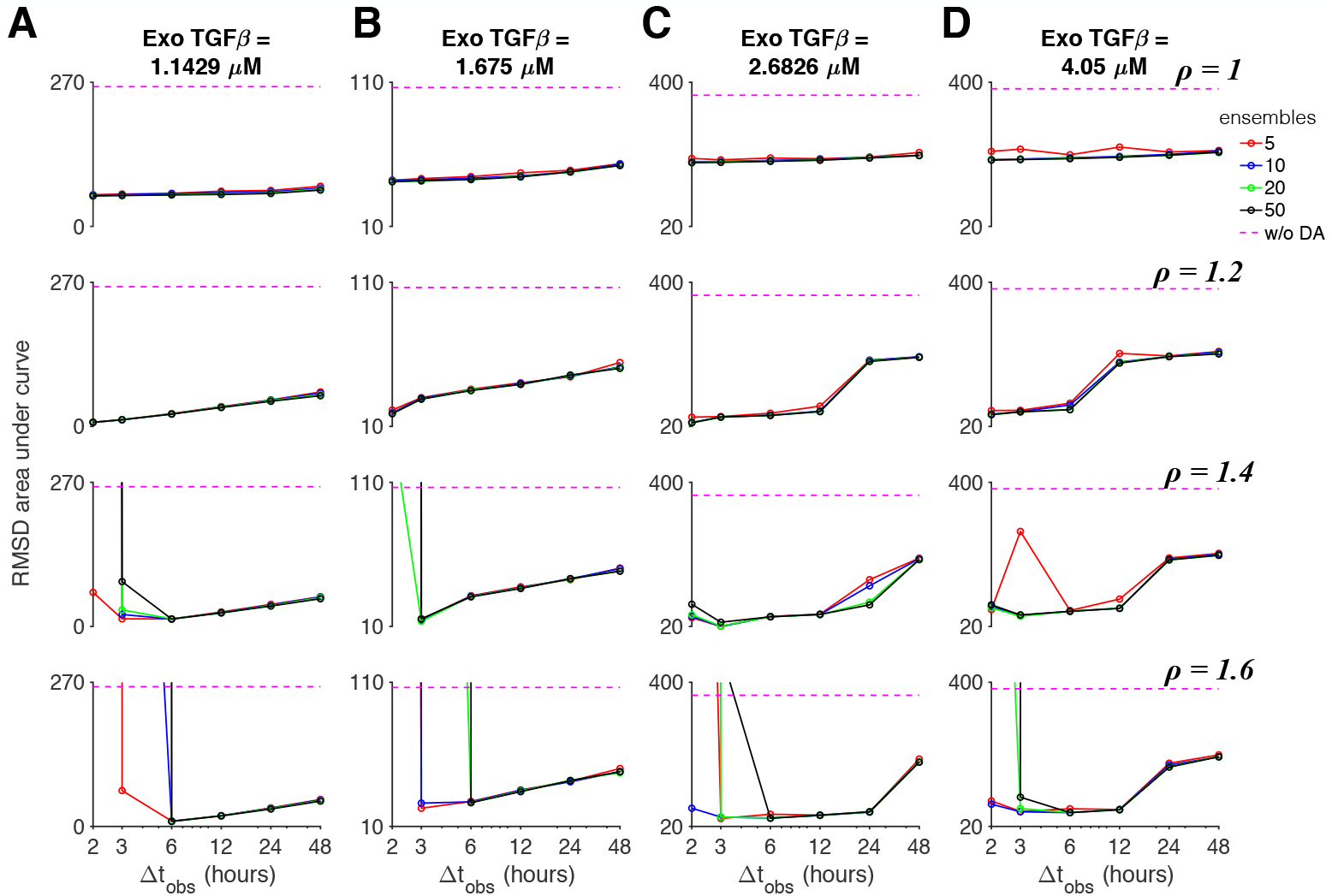
Optimal observation intervals and multiplicative inflation reduce prediction error of modified EMT dynamics, with model error. The area under the RMSD vs time curve is shown as a function of the observation interval Δ*t*_*obs*_, for different number of ensembles *k* (solid lines) and without data assimilation (DA, dashed magenta), for (A-D) four TGF*β* doses and multiplicative inflation values *ρ* (different rows). Parameters: Truth system: *k*_*d,s*_ = 0.108, Ensembles: *k*_*d,s*_ = 0.09.

## Discussion

In this study, we used a data assimilation approach on a series of synthetic experiments to forecast cell fate in the setting of epithelial-mesenchymal transition or EMT. First, proof-of-concept *in silico* experiments were performed in which experimental observations were produced from a known computational model with the addition of noise, but in the setting of no model error, i.e., all model parameters were assumed to be known. In the absence of model error, EMT dynamics were successfully reconstructed generally within 48 hours of observations.

To mimic parameter uncertainty present in *in vitro* experiments, we introduce model error in a manner which shifts the TGF*β* doses associated with state transitions. In the presence of model error, EMT dynamics were successfully reconstructed using the data assimilation approach incorporating multiplicative inflation and an optimal observation interval. That is, sufficiently frequent observation were needed to observe and predict EMT transitions, while a sufficient interval between observations and the addition of multiplicative inflation mitigate overconfidence in model predictions. With these ideal conditions, even in the presence of model error, the timing of EMT state transitions and steady state behavior were successfully predicted. Further, we found that these results negligibly depend on the number of ensembles in the EnKF, demonstrating that a computationally efficient approach using fewer ensembles is feasible and sufficient.

EMT is a process characterized by a phenotypic shift in epithelial cells to motile and oftentimes invasive mesenchymal cells. This tightly regulated process is fundamental in the generation of new tissues and organs during embryogenesis and is a key factor in tissue remodeling and wound healing [1–3]. While EMT is critical for development, its misregulation is implicated in many diseases, including cardiac fibrosis, cirrhosis, and cancer. In these disease states, it is not only crucial to understand what drives EMT in order to better understand the pathology, it is equally important to predict the timing of EMT-associated state transitions, with an eye towards developing effective therapies to reverse EMT-related disorders. One of the main complications with making such predictions is the limited number of EMT-associated markers that can be observed experimentally in any specific experiment; all experimental measurements are inherently providing an incomplete snapshot of the system state at a given moment in time.

The data assimilation approach presented in this study demonstrates several key advances in the prediction of EMT dynamics: (1) Expression levels of unmeasured EMT-associated cell markers are accurately reconstructed and predicted, based on a single ratiometric measurement of two cell markers. While these predictions are inherently limited by the details of the biophysical model from which they are based, this approach can be easily adapted to utilize more detailed predictive models of cell signaling to predict expression levels of additional unmeasured markers. (2) By integrating a predictive biophysical model with experimental observations, we can accurately predict future events, specifically the timing of cell phenotype state transitions. This technique can be more generally applied as a tool to probe responses to various experimental perturbations applied at different stages and timings throughout the EMT process, such as changes in TGF*β* dose or agonists and antagonists of different signaling pathways. Both of these extensions are the focus of ongoing future work. More broadly, potential future work will also explore the data assimilation performance in the setting of larger model error, accounting for potentially significant parameter uncertainty and differences between *in vitro* and *in silico* experiments.

A few prior studies have applied data assimilation approaches to different biological systems. Several studies, including those by two of the authors of this work, have reconstructed excitable cell dynamics, for various levels of scale and complexity. Munoz and Otani applied a Kalman filter on single cardiac cells to predict the dynamical behavior of state variables not directly observed, such as intracellular ionic concentrations [36]. Hoffman and colleagues used an ensemble Kalman filter approach to reconstruct complex electrical rhythms in one-dimensional and three-dimensional cardiac tissues, and similarly find that the addition of inflation in the data assimilation algorithm was pivotal to improve prediction accuracy, while also showing minimal influence on ensemble size [26, 27]. Several studies by Hamilton and colleagues have applied data assimilation approaches to predict dynamics of neural electrical activity, including determination of neural network connectivity [21], and reconstruction of intracellular ion concentrations [22] and of intracellular potential [23]. Moye and Diekman apply two different classes of data assimilation approaches to improve estimates of both neural cell state and model parameters for different types of bifurcation behavior [28]. Ullah and Schiff applied Kalman filters to predict unobserved states in neurons and small neural networks [24, 25].

Data assimilation has also successfully been applied to improve predictions of a human brain tumor growth in *in silico* experiments using synthetic magnetic resonance images [20]. Using predictions from a simple tumor growth model and integrating measurements from a more detailed model, the data assimilation algorithm successfully produced accurate qualitative and quantitative analysis of brain tumor growth. A similar Kalman filter approach has also been applied to dynamical state reconstruction, with a focus on prediction of unobserved state variables, and parameter estimation in a model of mammalian sleep dynamics [30] and blood glucose levels [29]. In general, prior work has focused on reconstructing physiological system dynamics, often predictions of unobserved system states, with several applications to excitable cells and tissue. While the dynamics of these systems are often governed by excitable, oscillatory, and bursting behavior, here we consider a system with distinct dynamics that are regulated by multiple bistable switches, and we show that data assimilation can successfully reconstruct cell state dynamics and transitions in such a system that governs cell phenotype.

As this study is an initial proof-of-concept demonstration of using data assimilation to predict EMT dynamics, there are several key limitations to be addressed in future studies. The Tian et al model used in this study represents the core regulatory pathway of TGF*β*-induced EMT. While the model is based on key experimental findings of the interactions of critical transcription factors and microRNAs regulating the EMT process [12], there are other signaling pathways, e.g., Wnt and *β*-catenin signaling [37, 38], involved in EMT that are not accounted for. However, our approach can be naturally extended to account for the details of additional signaling pathways. As an initial test, we only consider signaling occurring in a single cell and do not consider spatial interactions occurring within a multicellular tissue during EMT. Model development of spatial interactions during the EMT process is complex, and this challenge is indeed an area of ongoing work within our lab and others [39–41]. As described by Hunt and colleagues [16], the EnKF can be further extended to account for spatial localization and interacting spatial dynamics, and we plan to extend the approach demonstrated here to multicellular tissues in the future as well.

Additionally, we consider model error in the setting of a single inaccurate parameter, while multiple parameters are likely to be unknown or inaccurate in a more realistic scenario. However, here we consider the case in which model error arises due to differences in a single parameter, such that the source of model behavior is unambiguous and can be clearly attributed. Nevertheless, as noted above, we plan to determine how generalizable our results are by considering additional sources of model error in future work. Finally, our long-term goal is to consider realistic biological model error, that is, using our approach with in vitro observations from fluorescence measurements of the E-cadherin-ZEB dual sensor, and ultimately to predict cell fate during EMT.

## Supporting information

**S1 Table. Model state variables.** Notation, description, and epithelial-state initiation conditions for all state variables in the Tian et al model [12].

**S2 Table. Model parameters.** Parameters, description, baseline values, and units for the Tian et al model [12]. *Value unless otherwise noted.

## Acknowledgments

This work was supported through funding from the National Institutes of Health/National Institute of General Medical Sciences R01GM122855 (SHW, CAL) and the National Science Foundation DCSD-1762803 (EMC, MJH).

